# miRinGO: Prediction of biological processes indirectly targeted by human microRNAs

**DOI:** 10.1101/2020.07.24.220335

**Authors:** Mohammed Sayed, Juw Won Park

## Abstract

MicroRNAs are small non-coding RNAs that are known for their role in post-transcriptional regulation of target genes. Typically, their functions are predicted by first identifying their target genes and then finding biological processes enriched in these targets. Current tools for miRNA functional analysis use only genes with physical binding sites as their targets and exclude other genes that are indirectly targeted transcriptionally through transcription factors. Here, we introduce a method to predict gene ontology (GO) annotations indirectly targeted by microRNAs. The proposed method resulted in better performance in predicting known miRNA-GO term associations compared to the canonical approach. To facilitate miRNA GO enrichment analysis, we developed an R Shiny application, miRinGO, that is freely available from GitHub at https://github.com/Fadeel/miRinGO

## Introduction

Recent studies have suggested that microRNAs (miRNAs) are involved in many diverse biological processes and pathways including normal development and diseases [1]. Animal miRNAs bind to 3’UTR of mRNAs mainly through short sequence (6-8 NTs) called seed region and act as repressors of target gene expression [2]. Taking into account this short sequence binding, one miRNA can target hundreds or even thousands of genes and subsequently perturb many biological pathways [3].

In order to computationally predict miRNA-targeted pathways, typically potential target genes are compiled using one or more miRNA target prediction tools and a standard gene enrichment analysis [4] is used to find potential enriched pathways or gene ontology (GO) terms. Although the conventional pipeline is widely used, it has some limitations. Of these, existing tools consider only direct targets (post-transcriptionally regulated) of miRNAs but do not consider indirect targets (transcriptionally regulated) that are not necessarily enriched in miRNA seed-binding sites [5]. Indirect target genes are mainly regulated transcriptionally through transcription factors (TFs) [6, 7]. Furthermore, a study suggested that TFs are preferentially targeted by miRNAs [8].

Figure 1 shows a scenario where biological pathway/process can be missed by classical miRNA pathway analysis tools. Using the classical method, the percentage of targeted genes is (11%), on the other hand, if we include TF targets (i.e. indirect targets), the percentage of targeted genes will be (67%).

**Figure 1:**
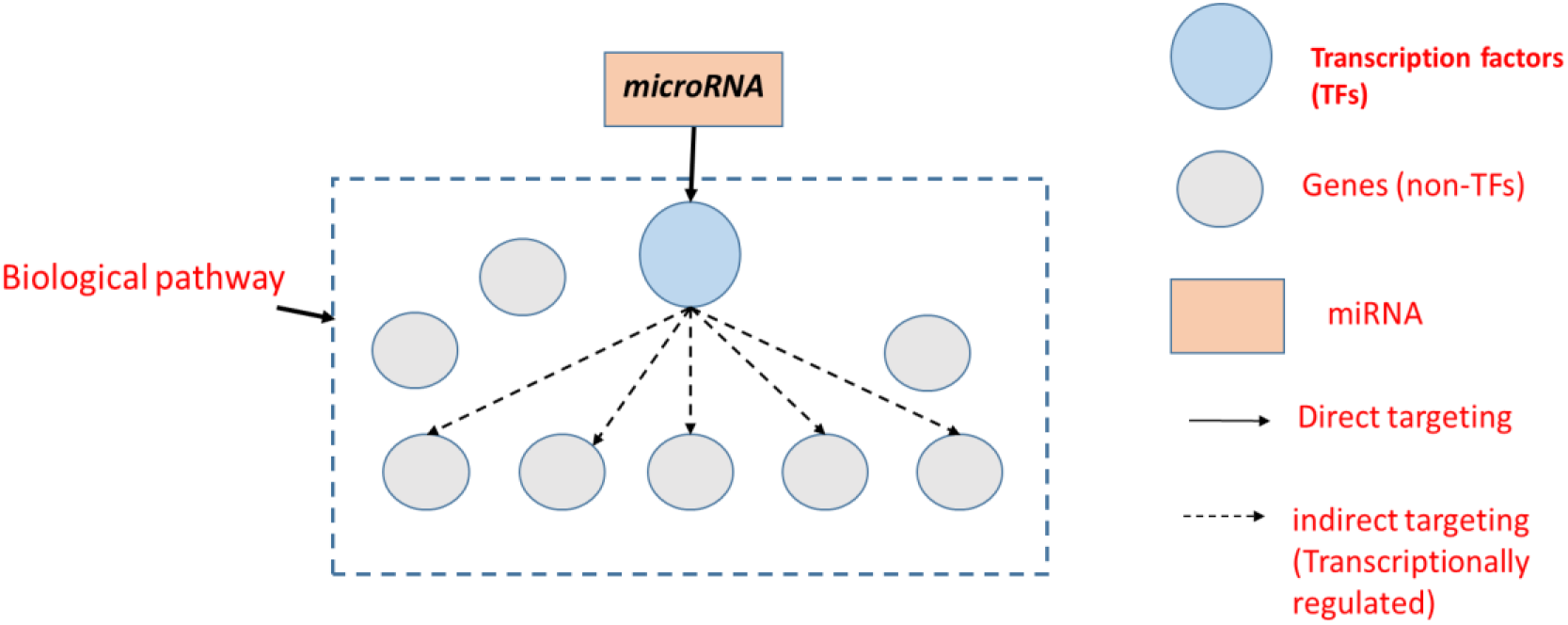
miRNAs can indirectly target biological pathways through transcriptions factors.

Indirectly regulated genes can be enriched in a specific biological pathway or phenotype. One notable example is the role of miR-200 family and miR-205 in controlling epithelial to mesenchymal transition (EMT) pathway through targeting *ZEB1* and *ZEB2* transcription factors [9]. Other studies have shown how miRNAs regulate cell differentiation through targeting TFs. Of these, Tay et al [10] demonstrated the role of miR-134 in embryonic stem cell differentiation through targeting *Nanog* and *LRH1*. Another study showed that miR-143 and miR-145 can work together to regulate smooth muscle cell differentiation and proliferation through targeting *Klf4* and *ELK1* transcription factors [11].

Several tools and web servers have been developed to predict potential biological pathways targeted by miRNAs [12–16]. While they are all similar in terms of using direct targets only, they use different databases for both miRNA targets and gene ontology annotations [17–21]. miTALOS [13] is the only tool that filters potential targets by incorporating tissue-specific genes. All tools except for StarBase [14] accept multiple miRNAs as input. A comprehensive comparison of widely used miRNA pathway analysis tools is shown in Table 1. In this study, we introduce miRinGO (<u>miR</u>NA <u>in</u>direct target <u>G</u>ene <u>O</u>ntology) that uncovers potential biological pathways affected by indirect targets of human miRNAs especially ones related to cell differentiation and development.

**Table 1:** A comparison of current tools of miRNAs pathway analysis

| Features/Tools | mirPath v3.0 | StarBase | miTALOS | miRWalk v3.0 |
| --- | --- | --- | --- | --- |
| Predicted targets databases | TargetScan (v6) / microT-CDS (v5.0) | TargetScan / miRanda/ PITA/ RNA22/ PicTar/... | TargetScan/ miRanda | TargetScan (v7.1) / miRDB |
| Validated targets databases | TarBase v7.0 | CLIP-Seq data | CLIP-Seq data | miRTarbase |
| Pathways / GO terms databases | KEGG/GO categories | KEGG/GO/ Reactome/ BioCarta | KEGG/ WikiPathways/ Reactome | KEGG/GO/ Reactome |
| **Inclusion of indirect targets?** | **No** | **No** | **No** | **No** |
| Tissue specific? | No | No | Yes | No |
| Allows multiple miRNAs? | Yes | No | Yes | Yes |

## Methods

### Overall pipeline

Our pipeline to predict indirectly-targeted biological processes by miRNAs consists of three steps as depicted in Figure 2. First, for each miRNA, potential directly-targeted TFs were compiled from the TargetScan database v7.2 [22]. Second, computationally predicted tissue-specific TF-gene associations were collected from the resources website of (Sonawane et al., 2017) [23]. In case of multiple input miRNAs, we use the intersection of indirect targets of each miRNA. Third, a hypergeometric test is conducted to find potential targeted biological processes.

**Figure 2:**
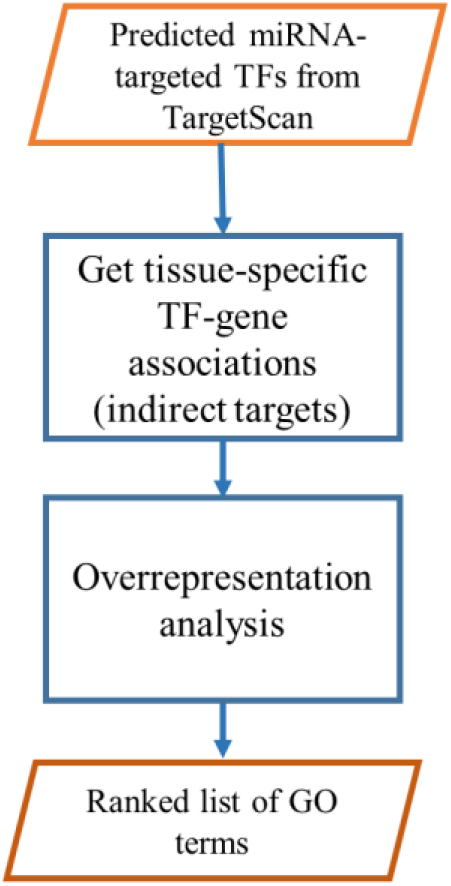
Pipeline of our miRNA GO enrichment analysis tool.

### Input Data

Data used by our tool was compiled from publicly available databases. Putative miRNA targets were downloaded from TargetScan v7.2 [22]. We downloaded the file with all predictions regardless of conservation of miRNA family or miRNA binding sites and kept high-confidence human miRNA targets (*Cumulative weighted context++ score* < −0.1).

Computationally predicted tissue-specific TF targets were downloaded from the resources website of (Sonawane et al., 2017) [23]. These TF targets were predicted using the PANDA (Passing Attributes between Networks for Data Assimilation) algorithm [24]. PANDA integrates three complimentary source of information i.e. TF sequences motif data, protein-protein interactions of TFs and gene co-expression from Genotype-Tissue Expression (GTEx) RNA-Seq data [25]. It contains TF-gene associations from 38 different tissues/tissue locations. We aggregated TF-gene associations from different locations but belong to the same tissue. We had 29 broad tissues after aggregation.

Gene ontology annotations were downloaded using Ensembl Biomart [26] (version GRCh38). GO terms with less than five genes were removed.

### Test dataset

To validate our method, we used a ‘gold standard’ dataset of miRNAs and their experimentally validated functions (GO terms) [27] from ftp://ftp.ebi.ac.uk/pub/databases/GO/goa/HUMAN/goa_human_rna.gaf. We filtered the dataset to include only high confidence annotations (excluded annotations with “*Inferred from Sequence or structural Similarity*” (ISS), “*Non-traceable Author Statement*” (NAS), and “*Traceable Author Statement*” (TAS) evidence codes). We also removed annotations with no reference article. To keep only relevant annotations, we removed generic GO terms shared by most miRNAs (e.g. “*miRNA mediated inhibition of translation*”, “*gene silencing by miRNA*” and “*gene silencing by RNA*”). Cell/tissue ontology was downloaded from http://www.ontobee.org/listTerms/CL?format=tsv. GO terms with less than five genes were removed. The filtered dataset consists of 335 pairs of miRNAs and their associated GO terms and is available in Supplementary Table S1.

## Results

### MicroRNA indirect vs direct targeting

To test the ability of our methodology to predict functions associated with a miRNA, we used a dataset with 335 known miRNA-GO term pairs. All TargetScan-predicted targets were included in this analysis. For each miRNA-GO term pair, resulting GO terms were ranked by the hypergeometric test *p-value* in ascending order, then rank values were converted to a percentile rank by dividing by the total number of GO terms. Finally, we picked the related GO term with the smallest *p-value* (smallest rank value). Known GO terms predicted by indirect targeting method have a significantly lower (Wilcox signed-rank test, one-sided *p-value* = 0.002417) rank compared to canonical direct targeting as shown in Figure 3.

**Figure 3:**
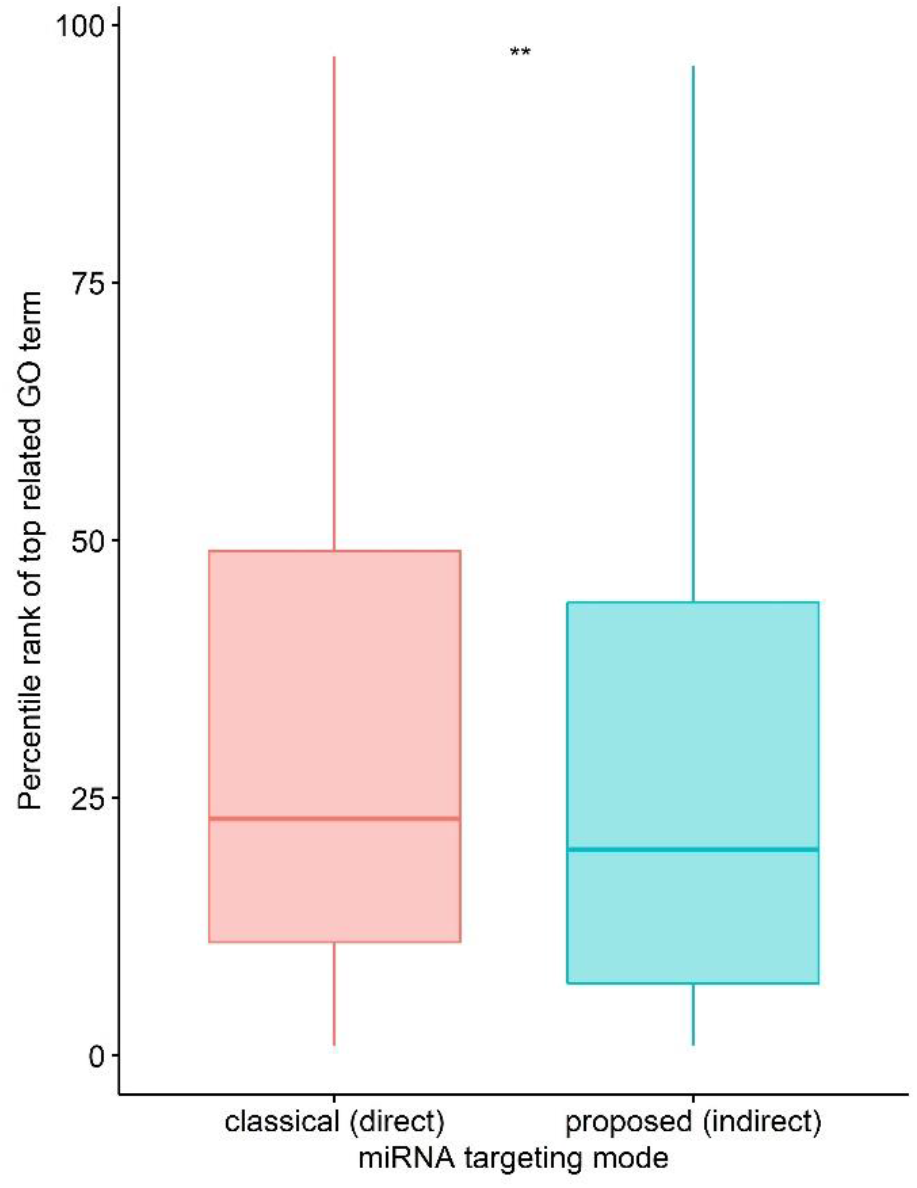
Comparison of indirect targeting with direct targeting (** represents p-value < 0.01)

### Effect of number of miRNA targets

Since miRNA GO enrichment analysis is affected by targets of the miRNA and TargetScan-predicted targets can have false-positives, we investigated the effect of the number of predicted miRNA targets on predicting the known GO terms. We repeated the same analysis but instead of using all predicted targets, we used top (20%, 40%, 60%, 80% and 100%) of potential targets (sorted by TargetScan context++ score [22]). Figure 4 shows that in all cases, average percentile rank of GO terms predicted by IT methodology is lower than DT. Although increasing the number of miRNA targets yielded a lower average rank (better performance), using all of targets did not give significantly better results compared to using the top 80% of targets and 40% of targets in case of indirect and direct targeting respectively.

**Figure 4:**
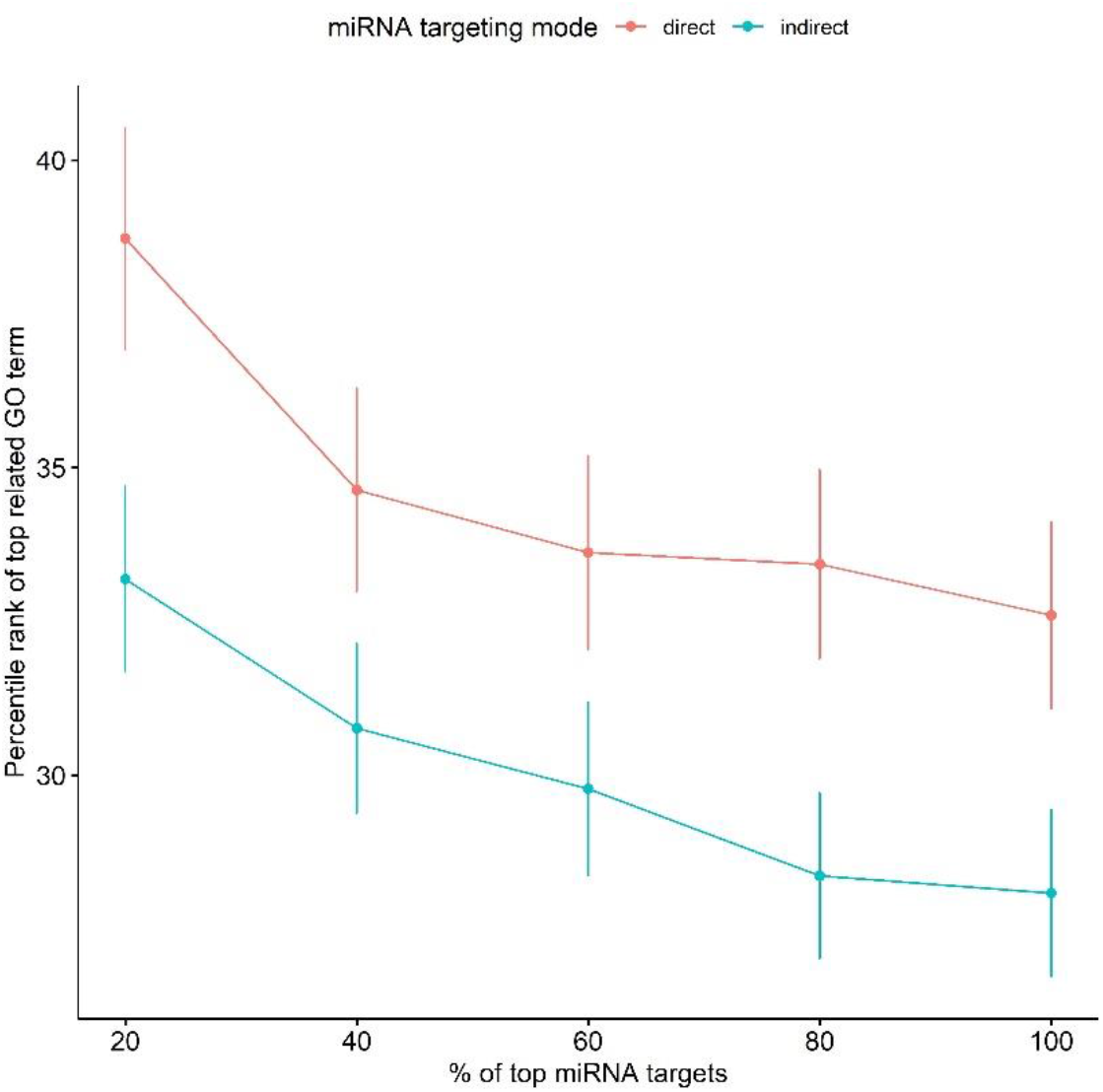
Effect of number of miRNA targets on miRNA GO enrichment analysis. Error bars represent one standard error.

### Indirect targeting reveals role of miRNAs in developmental processes

To investigate biological processes that are more likely to be affected by indirect targeting of miRNAs, we calculated *TF density* per GO term as defined by equation 1.

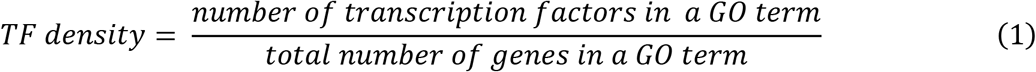

Table 2 shows top 5 GO BP terms with the highest TF density. All these GO terms are related to “*developmental process*” and all genes involved are transcription factors.

**Table 2:** Top 5 GO terms with the highest TF density.

| GO term ID | GO term | # of TFs | # of genes | Parent Process |
| --- | --- | --- | --- | --- |
| GO:0001714 | endodermal cell fate specification | 5 | 5 | developmental process |
| GO:0003211 | cardiac ventricle formation | 5 | 5 | developmental process |
| GO:0003357 | noradrenergic neuron differentiation | 5 | 5 | developmental process |
| GO:0021520 | spinal cord motor neuron cell fate specification | 7 | 7 | developmental process |
| GO:0021902 | commitment of neuronal cell to specific neuron type in forebrain | 7 | 7 | developmental process |

To see if transcription factors are enriched in development-related GO terms compared to other terms, we divided the GO terms (that have at least one TF) into two groups; one with development-related terms and the second with other terms or processes. We selected development-related terms by searching for GO biological process terms with the following keywords (“*development”, “cell fate”, “differentiation”, “stem cell”, “morphogenesis”, “cell specification”, “formation*”). Figure 5 shows that development-related terms (n = 613) tend to have significantly (*p-value* < 2.2e-16, Wilcoxon rank sum test) higher TF density compared to other terms (n = 1767).

**Figure 5:**
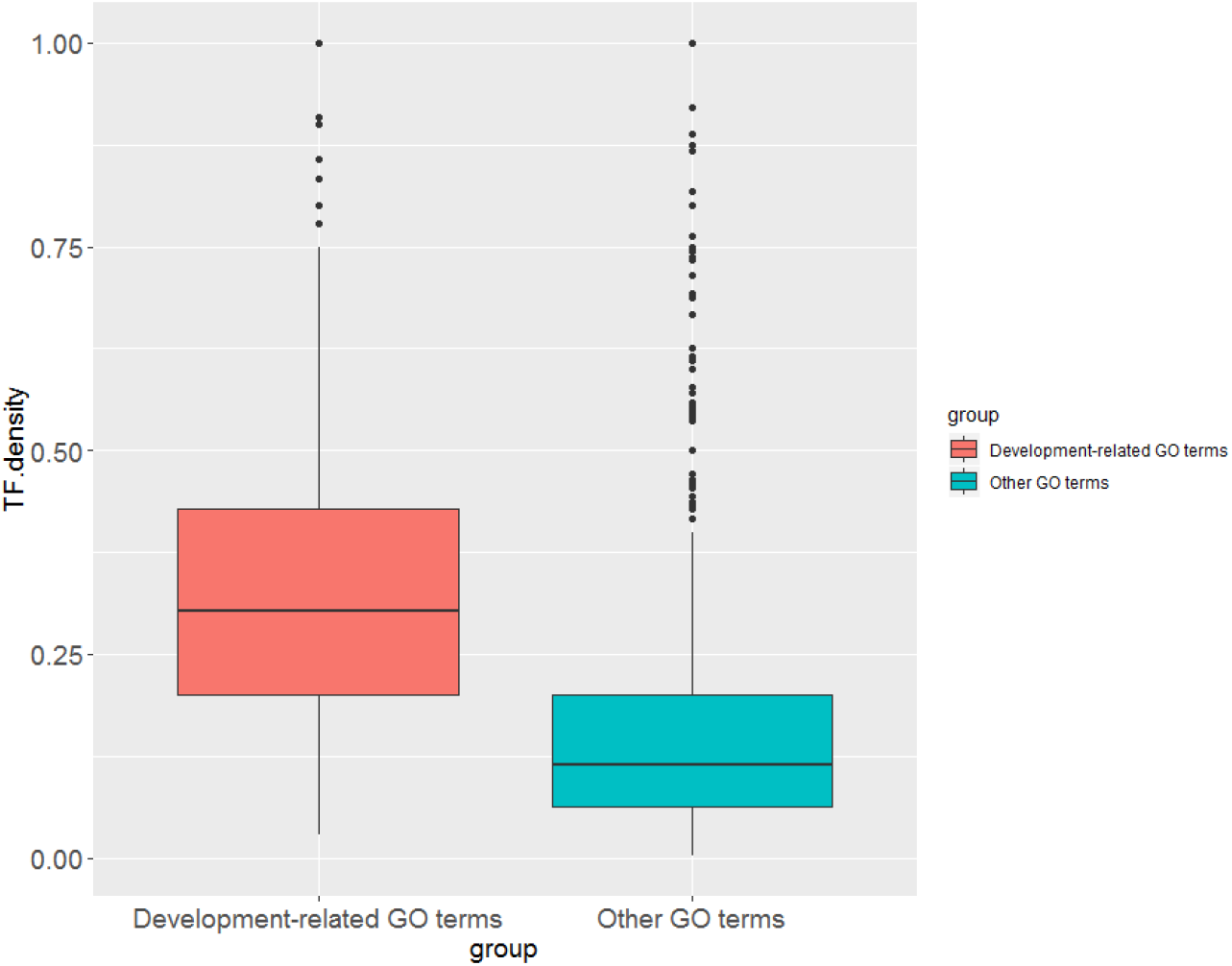
Comparison of TF density in development-related GO terms vs. all other terms Case Study: role of miR-9 in Neurogenesis.

To test the ability of our tool to capture relevant targeted development-related GO terms, we picked a miRNA with a known function to be able to compare our predicted GO terms with known ones. Of these miRNAs, miR-9 is brain-enriched miRNA and has prominent role in neurogenesis [28–30]. We ran our tool with the following inputs, we selected “brain” as the tissue type, “biological process” as the GO category, “indirect” as the targeting mode and “100” as the percentage of miRNA targets. Two out of the top 5 GO terms predicted are related to neurogenesis (“*Nervous system development*” and “*brain development*”).

To compare our results with existing miRNA pathway analysis tools, we downloaded predicted GO biological process terms for “miR-9” from 4 different web servers (accessed January 15, 2019): mirPath v3.0 [12], StarBase (mirTarPathway module) v3.0 [14], miTALOS v2 [13] and miRWalk v3.0 [15]. We searched for the highest-ranking GO term related to neurogenesis as shown in Table 3. Our tool ranked neurogenesis-related GO terms higher than other tools.

**Table 3:** Comparison of highest-ranking GO terms related to neurogenesis from different miRNA GO enrichment tools.

| Tool | Highest ranking GO term related to Neurogenesis | Rank |
| --- | --- | --- |
| miRinGO | Nervous system development | 1 |
| mirPath v3 | regulation of neuron maturation | 11 |
| miRWalk v3 | axonogenesis | 13 |
| StarBase v3 | Neurogenesis | 23 |
| miTALOS v2 | N/A | N/A |

### Multiple miRNAs GO analysis

In all miRNA GO analyses so far, we used one miRNA as an input. Several studies have shown that miRNAs can work together to regulate certain targets and biological processes [31, 32]. Of these, Gregory et al. [9] showed that the miR-200 family and miR-205 together regulate epithelial to mesenchymal transition (EMT). The miR-200 family consists of miRNAs with two different seed sequences: miR-200a/miR-141 and miR-200b/miR-200c/miR-429. We ran our tool with the following inputs: “kidney” as the tissue type, “biological process” as the GO category, and “indirect” as the targeting mode and “100” as the percentage of miRNA targets. The rank of *“epithelial to mesenchymal transition”* GO term (GO: 0001837) was lower when we used the intersection of indirect targets of these three miRNAs compared to ranks of GO terms predicted by each miRNA indirect targets as shown in Table 4.

**Table 4:** Effect of using multiple miRNAs in capturing EMT-related GO terms.

| miRNAs | Rank of top GO term related to EMT |
| --- | --- |
| miR-200a/miR-141 | 132 |
| miR-200b/miR-200c/miR-429 | 147 |
| miR-205-5p | 105 |
| All three miRNAs | 70 |

### R Shiny App

For ease of use of our method, we developed web app, miRinGO, using R Shiny package [33]. It has two panels, left one for input data and parameters selection and the right one for displaying the results in table format as shown in Figure 6. The tool provides users with the ability to choose different input data and parameters as detailed in Supplementary Table S2.

**Figure 6:**
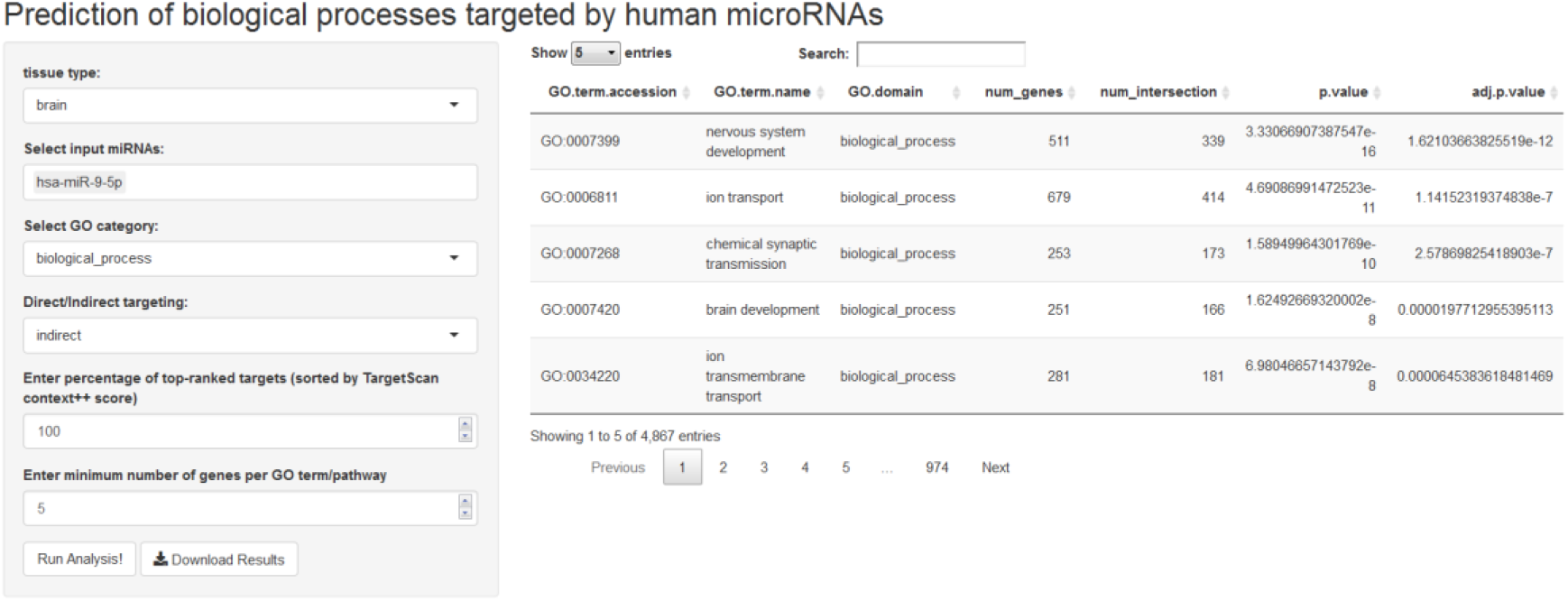
User interface of the R Shiny app, miRinGO.

## Discussion

We propose miRinGO a tool that detects biological processes indirectly targeted by miRNAs transcriptionally through transcription factors. Using miRinGO, we can include potential target genes even if there is no physical interaction between miRNA and the regulated genes. In order to validate this method, we used a dataset of miRNAs and their known targeted GO terms [27]. Although this dataset is considered a significant step towards having a gold standard to validate different miRNA pathway or GO analysis tools, it is still limited to a fraction of human miRNAs and focused more on cardiovascular-related processes. Using this dataset, however, indirect targeting showed better performance in predicting known targeted processes compared to direct targeting method, even if we use different fractions of input miRNA targets. It is also worth noting that although increasing the number of miRNA targets yielded better performance, using all of targets did not give significantly better results compared to using top 80% of targets and 40% of targets in case of indirect and direct targeting, respectively. This could be due to the fact that miRNA target prediction tools suffer from having many false positives [34].

Since our method is mainly focused on miRNA-targeted TFs and development-related GO terms or pathways have more TFs than other terms, it is more suitable to use this tool to uncover the tissue-specific roles of miRNAs in development and cell differentiation. Using this method, we predicted biological pathways known to be targeted by miR-9, a miRNA with a known role in neurogenesis. Tan et al. [35] showed that miR-9 regulates neural stem cell differentiation and proliferation by targeting *HES1* transcription factor. Using indirect targeting, three genes related to neuron differentiation (*FEZF2, SOX3* and *ZHX2*) that are predicted to be targeted by *HES1* (but are not direct targets of miR-9-5p) are now included in GO enrichment analysis as indirect targets of miR-9-5p.

One limitation of our method is that we use to two sets of computationally-predicted targets: one for miRNA direct targets and the other for tissue-specific TF targets. This might increase the effect of false positives in miRNA GO enrichment analysis. This limitation can be partly alleviated by using only high-confidence miRNA targets (i.e. ones with smaller TargetScan *context++* score). Although our method outperformed current miRNA GO analysis method, it is not intended to replace standard miRNA GO analysis method but on the other hand, to give a different perspective of miRNA roles in regulating biological processes and to uncover ones that are previously overlooked by current tools, especially ones related to development and cell differentiation.

## Notes

### Competing Interest Statement

The authors have declared no competing interest.

